# ADHD-like traits reshape the balance between inhibitory control and predictive processes

**DOI:** 10.1101/2025.10.28.685045

**Authors:** Karolina Horváth, Bianka Brezóczki, Adrienn Holczer, Teodóra Vékony, Dezső Németh

**Affiliations:** Gran Canaria Cognitive Research Center, Atlántico Medio University, Las Palmas de Gran Canaria, Spain; Doctoral School of Psychology, ELTE Eötvös Loránd University, Budapest, Hungary; Institute of Psychology, ELTE Eötvös Loránd University, Budapest, Hungary; Brain, Memory and Language Research Group, Institute of Cognitive Neuroscience and Psychology, HUN-REN Research Centre for Natural Sciences, Budapest, Hungary; BML-NAP Research Group, Institute of Psychology, Eötvös Loránd University and Institute of Cognitive Neuroscience and Psychology, HUN-REN Research Centre for Natural Sciences, Budapest, Hungary; Centre de Recherche en Neurosciences de Lyon CRNL U1028 UMR5292, INSERM, CRNS, Université Claude Bernard Lyon 1, 69500 Bron, France

**Author notes:** Correspondence: Correspondence should be addressed to Dezso Nemeth at dezso.nemeth [at]inserm.fr. These authors contributed equally to this work (Karolina Horváth, Bianka Brezóczki, Teodóra Vékony, Dezső Németh).

**Keywords:** Attention Deficit/ Hyperactivity Disorder, statistical learning, cognitive control, cognitive tradeoff, spectrum-approach

## Abstract

Adaptive behavior relies on a dynamic balance between flexible, goal-directed control and efficient, automatic processes. This equilibrium is often disrupted in Attention-Deficit/Hyperactivity Disorder (ADHD), a condition understood to exist along a continuum of traits in the general population. While ADHD is consistently linked to response inhibition deficits, its relationship with statistical learning (a mechanism for habit learning) and, crucially, the interaction between these functions, remains underexplored. Here, using a novel paradigm in a non-clinical sample of university students (n = 226), we investigated how ADHD-like traits modulate the interplay between response inhibition and statistical learning. We found that higher ADHD-like traits were associated with poorer response inhibition, confirming previous research. Importantly, we uncovered an antagonistic relationship between inhibition and statistical learning, where weaker inhibitory control typically led to enhanced learning of environmental regularities. However, this learning advantage progressively diminished across the ADHD trait continuum, becoming markedly reduced in individuals with high symptom prevalence. These findings provide novel evidence that ADHD-like traits influence not only isolated neurocognitive processes but also their dynamic interaction, highlighting a spectrum-based mechanism that may underlie the transition from adaptive variability to maladaptive behavioral patterns. This work advocates for a dimensional approach to ADHD, emphasizing early detection and targeted interventions for individuals with varying levels of symptomatic expression, broadening support beyond traditional diagnostic boundaries.

## Introduction

Navigating a complex and ever-changing world requires a dynamic interplay between flexible, goal-directed control and efficient, automatic processes. The capacity for goal-directed behavior allows for deliberate adaptation to novel demands, albeit at a significant cognitive cost (Grahek et al., 2023; Kurzban et al., 2013; Musslick & Cohen, 2021), while automatic processes enable rapid and efficient execution of well-learned actions at the expense of flexibility (Balleine & O’Doherty, 2010). An optimal balance between these two systems is fundamental for adaptive functioning. However, this equilibrium can be disrupted by structural or functional alterations within these neurocognitive systems (Gillanet al., 2015, 2016). Attention-Deficit/Hyperactivity Disorder (ADHD) is a prime example of such a disruption, characterized by enduring patterns of inattention and hyperactivity-impulsivity (American Psychiatric Association, 2022). A prevailing hypothesis posits that impaired goal-directed processes are a core neurocognitive deficit in ADHD (Barkley, 1997; Bush et al., 2005; Castellanos et al., 2006; Häger et al., 2021; Martel et al., 2007; Molitor et al., 2019; Natsheh et al., 2019; Seidman, 2006; Silverstein et al., 2020), which consequently perturbs the typical balance between goal-directed and automatic mechanisms (Hogarth et al., 2012). While the clinical presentation of ADHD is well-defined, mounting evidence suggests it is not a categorical disorder but rather exists on a continuum, with traits of inattention and hyperactivity-impulsivity distributed across the general population (Greven et al., 2016; Hengartner & Lehmann, 2017; Herrmann et al., 2009; Levy et al., 1997; T. Li et al., 2019; Paloyelis et al., 2009; Panagiotidi et al., 2018, 2019; Polner et al., 2015; Schippers et al., 2024; Vogel et al., 2018). This dimensional perspective provides a powerful framework to investigate how subtle variations in ADHD-like traits modulate the fundamental trade-off between cognitive control and automaticity in a non-clinical population.

To investigate this trade-off along the non-clinical ADHD spectrum, it is essential to examine the core cognitive functions involved. Goal-directed behavior depends on prefrontal cortex–related executive functions (Diamond, 2013; Friedman & Robbins, 2022; Miyake et al., 2000), such as response inhibition, cognitive flexibility and working memory (Chambers et al., 2009; Wright et al., 2014). Prior research on isolated components of executive functioning has predominantly focused on response inhibition (Crosbie et al., 2013; Hadas et al., 2021; Hart et al., 2014; Li et al., 2025; Ma et al., 2012; Nigg, 2001; Nigg et al., 2002; Silverstein et al., 2020; Wodka et al., 2007). Response inhibition is an ability to suppress inappropriate or prepotent actions (Chambers et al., 2009; Wright et al., 2014). A substantial body of evidence indicates that impairments in response inhibition are present in ADHD, both structurally and functionally (Bush et al., 2005; Castellanos & Tannock, 2002; Crosbie et al., 2013; Hadas et al., 2021; Hart et al., 2014; X. Li et al., 2025; Liston et al., 2011; Ma et al., 2012; Nigg, 2001; Nigg et al., 2002; Silverstein et al., 2020; Wodka et al., 2007). These alterations have been observed not only in clinical populations but also in individuals with non-clinical ADHD-like traits, where such traits are associated with reduced response inhibition performance and, consequently, greater impulsive behavior (Crosbie et al., 2013; Polner et al., 2015). In the context of balancing goal-directed and automatic processes, diminished inhibitory control may lead to a greater reliance on the automatic system, including mechanisms such as statistical learning (SL) (Nemeth et al., 2013). This is supported by the competition hypothesis, which posits an antagonistic relationship between executive control and statistical learning. Executive functions rely on voluntary, resource-demanding mechanisms mediated by a network involving the medial temporal lobe and the prefrontal cortex (Friedman & Robbins, 2022; Miyake et al., 2000; Nemeth et al., 2013). Because the prefrontal cortex is responsible for response inhibition, its diminished activity reduces top-down interference. In contrast, automatic processes are mediated largely by the striatum, which facilitates nonconscious, SL (Poldrack et al.; 2011; Turk-Browne et al.; 2010). Extensive evidence suggests that these two systems compete for cognitive resources (Daw et al.; Watson et al., 2018). Specifically, suppressing the explicit, hypothesis-testing system can lead to preserved or even enhanced striatum-dependent learning (Brown & Robertson, 2007; Foerde et al., 2006; Fu & Anderson, 2008). Conversely, increasing the reliance on attention-dependent, explicit processes has been shown to impair SL (Howard & Howard, 2001). Thus, when inhibitory control is weakened (as seen in individuals with higher ADHD-like traits) the automatic system may function more efficiently due to a lack of competition from executive systems. Within this framework, SL is defined as the unconscious acquisition of environmental regularities that support predictive processing and underlie the development of habits and skills (Aslin, 2017; Conway, 2020; Horváth et al., 2022; Theeuwes et al., 2022). While the majority of studies report largely intact SL in both clinical (Adi-Japha et al., 2011; Karatekin et al., 2009; Pedersen & Ohrmann, 2018; Takács et al., 2017; Vloet et al., 2010) (for exceptions, see [Barnes et al., 2010]), and non-clinical populations with ADHD-like traits (Parks & Stevenson, 2018), the interplay between inhibition and SL may still be significantly modulated by ADHD traits. We therefore posit that the interaction between these two distinct neurocognitive systems, impaired response inhibition and preserved SL, represents a key mechanism underlying the behavioral profile of ADHD-like traits on a continuum, a relationship that has yet to be systematically investigated.

Accordingly, the core limitation of prior research is that these processes have primarily been examined in isolation, whereas real-world learning and adaptive behavior typically rely on the coordinated operation of multiple neurocognitive functions (Horváth et al., 2022; Pedraza, Farkas, et al., 2024). As such, this isolated approach is limited, since dysfunctions may emerge when these processes are forced to interact (Balleine & O’Doherty, 2010; Castellanos & Tannock, 2002). To date, only a single study in a clinical child population has investigated such interactions, revealing a bias toward automatic behavior in ADHD alongside disrupted goal-directed control (Natsheh et al., 2019). Importantly, ADHD in childhood and adulthood may differ at the neurocognitive level due to the maturation of prefrontal regions (Shaw et al., 2007). Therefore, it is crucial to examine the interactive relationship between SL processes and goal-directed versus automatic behavior.

Moreover, to better capture the underlying mechanisms of ADHD symptomatology, it is essential to move beyond traditional dichotomous clinical–control group comparisons. In the present study, we adopt a spectrum-based methodological approach to examine how individual differences in ADHD-like traits relate to neurocognitive processes, specifically the interplay between inhibitory control and SL. Based on existing literature, we hypothesized an antagonistic relationship between SL and inhibitory control, such that reduced inhibitory control may facilitate stronger statistical learning (Brown & Robertson, 2007; Foerde et al., 2006; Fu & Anderson, 2008; Nemeth et al., 2013). Building on prior research, we further hypothesized that this relationship would be modulated by the prevalence of ADHD-like traits, with the interaction between SL and inhibitory control varying systematically across the trait spectrum rather than following a simple linear pattern (Ambrus et al., 2020; Horváth et al., 2022; Janacsek et al., 2012; Nemeth et al., 2013; Pedraza, Farkas, et al., 2024; Pedraza, Vékony, et al., 2024).

## Methods

### Participants

We recruited three hundred and seventeen university students who participated in an online experiment and received course credit for their participation. All participants provided written informed consent, and the research protocol adhered to ethical guidelines established by the Declaration of Helsinki. All protocols were approved by the Eötvös Loránd University Research Ethics Board (2024/214). The dataset for this study is the same as that used by Vékony et al. (2025).

Several exclusion criteria were applied to ensure good data quality. Participants were excluded if they reported any neurological or psychiatric conditions (*n* = 43), were currently taking medications known to influence central nervous system functioning (*n* = 8), had consumed alcohol within three hours of testing (*n* = 19), or possessed prior familiarity with ASRT paradigms (*n* = 10). Incomplete data also resulted in exclusion, particularly for participants who did not complete the Adult ADHD Self-Report Scale (ASRS; *n* = 25) or who failed the attention test during the questionnaires (*n* = 7). To ensure participants were engaged, a statement was included in the questionnaires: “If you are paying attention right now, please select the ‘Agree’ option.” Participants who did not follow this instruction were considered inattentive, and their data was excluded from the final analysis.

Task performance served as an additional screening criterion. Participants demonstrating accuracy rates below 80% (*n* = 11) were excluded to ensure adequate task engagement. Similarly, those requiring more than 50 minutes to complete the task (*n* = 7) or who restarted the procedure during testing (*n* = 3) were removed from the analyses. Following these exclusions, 91 participants were removed in total. Notably, some satisfied multiple exclusion criteria, resulting in a final sample of 226 participants (retention rate: 71.6%). The mean age of the participants was 21.42 ± 4.76 SD years (183 female, 42 male, and 1 non-specified). The sample of 226 participants showed a varied educational background: 178 participants (78.8%) had a high school education or lower, 41 (18.1%) had a bachelor’s degree or equivalent, and 7 (3.1%) had a master’s degree or equivalent.

### The Cognitive Trade-off Task (CTT)

The Cognitive Trade-off Task (CTT) integrates the Alternating Serial Reaction Time Task (ASRT), a well-established visuomotor statistical learning paradigm (Farkas et al., 2023; Howard & Howard, 1997; Janacsek et al., 2012), with a Go/No-Go component, traditionally employed to assess response inhibition. In the present study, we adapted the ASRT task to simultaneously measure statistical learning and inhibitory control by incorporating a Go/No-Go component.

The experimental task was programmed in JavaScript using the jsPsych library v.6.1.0. (Vékony, 2021). Depending on the experimental condition (detailed below), either a dog or a cat head appeared in one of four positions arranged horizontally in the center of the screen (Figure 1A). The four circles were positioned in the middle of the screen, and participants received instructions to press the designated keyboard button (’S’,’F’,’J’, or’L’ from left to right) as rapidly and accurately as possible when presented with go stimuli, corresponding to the stimulus position. Conversely, when presented with No-Go stimuli, participants were instructed to suppress their response and refrain from pressing any button.

**Figure 1.**
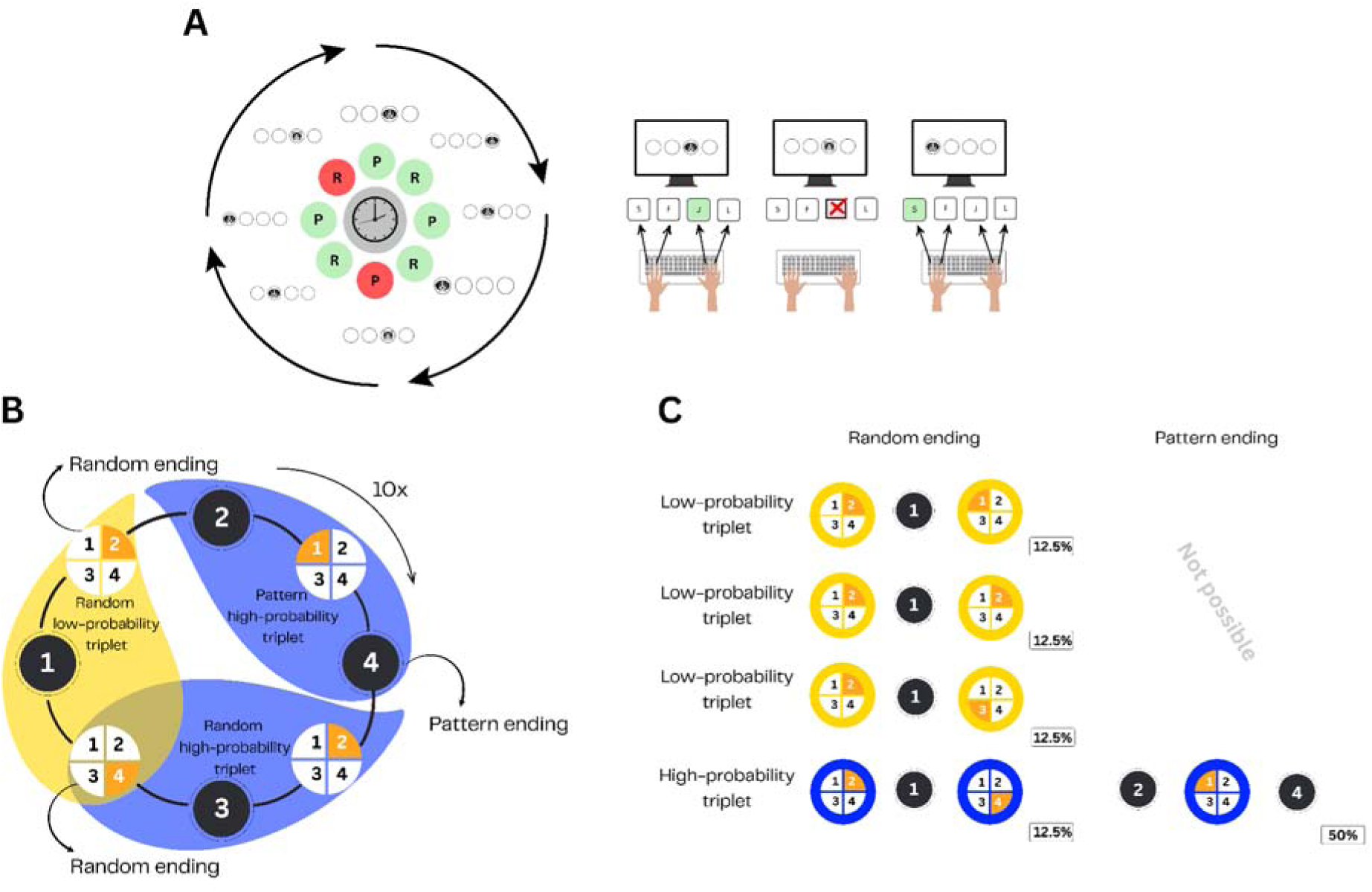
The Cognitive Trade-off Task. **(A)** Participants completed 30 blocks, each containing 80 trials. In each trial, a ‘go’ stimulus appeared on the screen, and participants were instructed to press the corresponding button (S, F, J, or L) as quickly and accurately as possible. Ten percent of the trials were ‘No-Go’ stimuli, which required participants to withhold their response. A hidden structure was embedded in the task, where every second trial followed a predetermined, 8-element probabilistic sequence. **(B)** The alternating structure of the task led to certain three-trial combinations (triplets) occurring more frequently than others. These were categorized as high-probability triplets, while the less frequent combinations were considered low-probability triplets. The elements within these triplets were either pattern elements (indicated by a black background) or random elements (indicated by an orange background). **(C)** Due to the alternating structure of the task, high-probability triplets were formed in two ways. The most common configuration, occurring in 50% of the trials, consisted of two pattern elements and one random element in the middle. The second configuration, occurring on 12.5% of the trials, was formed by two random elements with a pattern element in the middle. In total, 62.5% of trials were categorized as high-probability triplets. The remaining 37.5% of trials were considered low-probability triplets.

When participants executed the correct button press on’go’ trials, the target stimulus disappeared immediately, followed by a 120 ms interstimulus interval before the presentation of the subsequent stimulus. Incorrect button presses on ‘go’ trials resulted in the stimulus remaining visible until participants selected the correct response. Notably, in 10 out of every 80 trials, either a cat head or a dog head stimulus functioned as the ‘No-Go’ signal, requiring participants to suppress any button press. The No-Go stimuli remained displayed for 1000 ms before disappearing, regardless of the participants’ responses.

Unbeknownst to participants, an 8-element sequence was continuously repeated throughout the experimental session. Within the eight-element sequence, sequence components alternated with random elements. Consequently, the presentation of a sequence stimulus was invariably followed by a random element, creating a consistent alternating pattern (e.g., 2 – r – 4 – r – 3 – r – 1 – r, where’r’ represents a random location, while the numbers indicate the predetermined location of the sequence) (Figure 1B).

This alternating structure of the ASRT task led to certain three-trial combinations, known as triplets, occurring more frequently than others. Some triplets occur with a higher probability (high-probability triplets) than other triplets (low-probability triplets). Trial classification depends on the final trial within the triplet, independent of whether it constitutes a random trial. For example, in the sequence 2 – r – 4 – r – 3 – r – 1 – r, high-probability triplets are, e.g., 2-r-4 and 4-r-3. Conversely, using the same sequence, low-probability triplets are, for example, 2-r-3 and 3-r-4. During the experiment, high-probability triplets were constructed either with two pattern trials and one random trial in the middle (50% of cases) or with two random trials and one pattern trial in the middle of the triplet (12.5% of trials). All other triplets were defined as low-probability triplets, which appeared in 37.5% of the trials. In total, 62.5% of trials functioned as the concluding element of high-probability triplets, while 37.5% of trials served as the final element of low-probability triplets (Figure 1C).

The experimental design utilized counterbalanced stimulus assignments to mitigate potential animal (cat or dog) preference effects. Half of the participants were assigned to respond to a dog head (go condition) while inhibiting responses to a cat head (No-Go condition), whereas the remaining participants received the opposite stimulus-response mapping. The appearance of’No-Go’ stimuli was randomized across blocks, appearing at both pattern and random positions within the alternating sequence. The structure also ensured that no two consecutive’No-Go’ stimuli were presented, and the initial two trials of each block were never’No-Go’ trials.

Participants completed 30 experimental blocks. Within each block, participants encountered the 8-element sequence 10 times, resulting in 80 stimuli per block. Each block contained 10 No-Go stimuli and 70 Go stimuli, maintaining a consistent 12.5% No-Go trial ratio throughout the experiment.

### Adult ADHD Self-Report Scale (ASRS v1.1; Kessler et al., 2005)

ADHD tendencies were assessed using a self-report questionnaire, which enabled us to identify the prevalence of ADHD-like symptoms along the spectrum. The Adult ADHD Self-Report Scale (ASRS) was selected to provide a quantitative assessment of ADHD symptom prevalence within our non-clinical sample (Kessler et al., 2005). This psychometric instrument represents a widely validated screening tool developed specifically for identifying ADHD symptoms in adult populations (Kessler et al., 2005). The ASRS comprises 18 self-report items utilizing a 5-point Likert scale format (ranging from 0 = never to 5 = very often) to evaluate symptom frequency over the preceding six-month period. The maximum possible score on the ASRS questionnaire is 72 points. In our study, the highest score was 57 (Figure 2). This bifactorial structure aligns with the DSM-5 conceptualization of ADHD symptom presentation (American Psychiatric Association, 2022). The total ASRS score for each participant was calculated and used as a measure of ADHD-like symptoms in our analyses.

**Figure 2.**
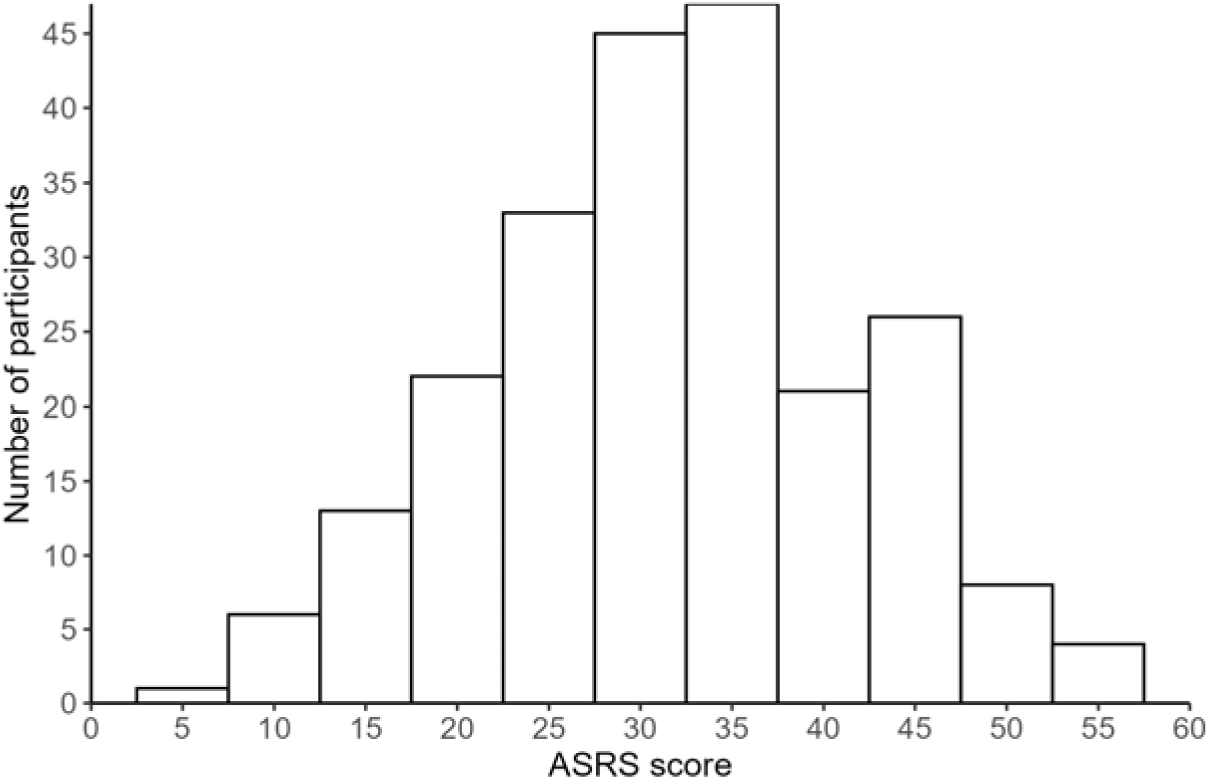
Distribution of the ASRS scores. The scores, which can range from 0 to 72, extended from a minimum of 6 to a maximum of 57 (M = 31.66, SD = 10.04). The score distribution exhibited a slight negative skewness (skewness =-0.013). The majority of participants scored in the mid-range of the scale.

### Procedure

The experiment was hosted by the Gorilla Experiment Builder platform (https://gorilla.sc; (Anwyl-Irvine et al., 2020). Upon initial access, participants reviewed and provided informed consent before proceeding to the experimental session. The testing session started with the CTT, which consisted of 30 blocks. Post-task questioning revealed that no participants could identify the hidden sequence or describe any underlying pattern, demonstrating that the task appropriately measured statistical learning (Horváth et al., 2022; Vékony et al., 2022). Additionally, demographic information was collected, including age, educational background, and other relevant participant characteristics. One week later, participants were asked to complete a battery of self-report questionnaires on Qualtrics, which included the ASRS.

### Data preparation

All data processing, statistical computations, and visualization were performed using R version 4.4.2. (R Core Team, 2024). Prior to analysis, trials were categorized based on whether they served as the final element of a high-probability or low-probability triplet. For the calculation of statistical learning measures,’No-Go’ trials were excluded. Trials with response times below 100 ms or above 1000 ms were removed. Furthermore, trials falling below or beyond three standard deviations were excluded to control for extreme outlier responses. Repetitions (e.g., 3-3-3) and trills (e.g., 2-1-2) were excluded because participants typically demonstrate faster responses to these sequences due to reduced motor complexity rather than statistical learning effects (Soetens et al., 2004). Trials with incorrect responses were excluded.

### No-Go performance score calculation

We measured response inhibition performance through’No-Go’ trials. Block wise accuracy on’No-Go’ trials was calculated as the proportion of trials where participants successfully inhibited their response. Correct inhibition was defined as refraining from pressing the corresponding key throughout the 1000 ms stimulus duration.

### Statistical analysis

We applied linear mixed-effects modeling (LMM) using the *mixed* function from the *afex* package (Singmann et al., 2024). We initially implemented the maximal random effect structure. If convergence issues or singular fit warnings occurred, we first removed correlations between random variables. If they persisted, we removed interactions between variables. We retained the most complex models that converged adequately. Post-hoc analyses of simple means and simple slopes were performed using estimated marginal means and estimated marginal trends calculated with the *emmeans* package (Lenth, 2018). Fixed effects were evaluated using Type III tests. For significant interactions involving continuous predictors, we divided continuous variables based on their mean values, with cut-points set at ±1 SD from the mean. This approach was applied when No-Go accuracy, Block, or ASRS scores were treated as continuous variables. Statistical significance was set at α = 0.05 for all tests. Data visualization was performed using the *ggplot2* package, with additional support from *ggplot* and *afex*.

We constructed two models to address our research questions. We initially specified the maximal random-effects structure to test our hypotheses, which was systematically simplified in cases of non-convergence until convergence was achieved. First, we examined how the level of ADHD-like traits related to response inhibition. Second, we investigated how ADHD-like traits and response inhibition interact to influence statistical learning. In our first model, the outcome variable was No-Go accuracy (measuring response inhibition; in percentage), calculated per participant for each block, with block (within-subject) and ASRS total score (between-subject) as predictor variables. Block and ASRS total scores were mean-centered. In our second model, the outcome variable was reaction time, calculated per participant for each Block and Triplet Type, with Block, Triplet type, and No-Go accuracy as within-subject predictors, and ASRS total score as a between-subject predictor. Block, No-Go accuracy, and ASRS total score were all mean-centered.

1. Model_1_: *No-Go accuracy ∼ Block x ASRS score + (Block | ID)*
2. Model_2:_ *Go RT ∼ Block x Triplet type x ASRS score x No-Go accuracy + (Block | ID)*

## Results

### Higher ADHD-like traits are associated with decreased response inhibition

We fitted a linear mixed-effects model with accuracy on No-Go trials as the outcome variable. This was operationalized as the mean percentage of correct’No-Go’ responses per block. Fixed effects included Block (centered), ASRS scores (centered), and their interaction. Both predictors were mean-centered and treated as continuous variables in the model. Random effects included participant ID and random slopes for block within participants, allowing individual variation in inhibitory control trajectories across time and estimating the correlation between participants’ starting levels and their rate of change. The model revealed a significant main effect for both Block (*F*_(1, 224.61)_ = 153.11, *p* <.001) and ASRS score (*F*_(1, 223.94)_ = 8.59, *p* =.004), indicating that response inhibition significantly differed across levels of both factors. Response inhibition was worse with higher levels of ASRS scores (*b* =-0.003, SE = 0.001) (Figure 3), and significantly decreased as the task progressed (*b* =-0.006, SE = 0.001). However, the interaction between ASRS scores and Block was not statistically significant (*F*_(1, 224.09)_ = 0.44, *p* =.509), suggesting that the effect of ASRS score on response inhibition did not change with the progress of the task. These findings revealed that higher ADHD-like traits were associated with a decrease in response inhibition. Full results are presented in Supplementary Table 1, including b, SE b, 95% CI, t, df, and p values.

**Figure 3.**
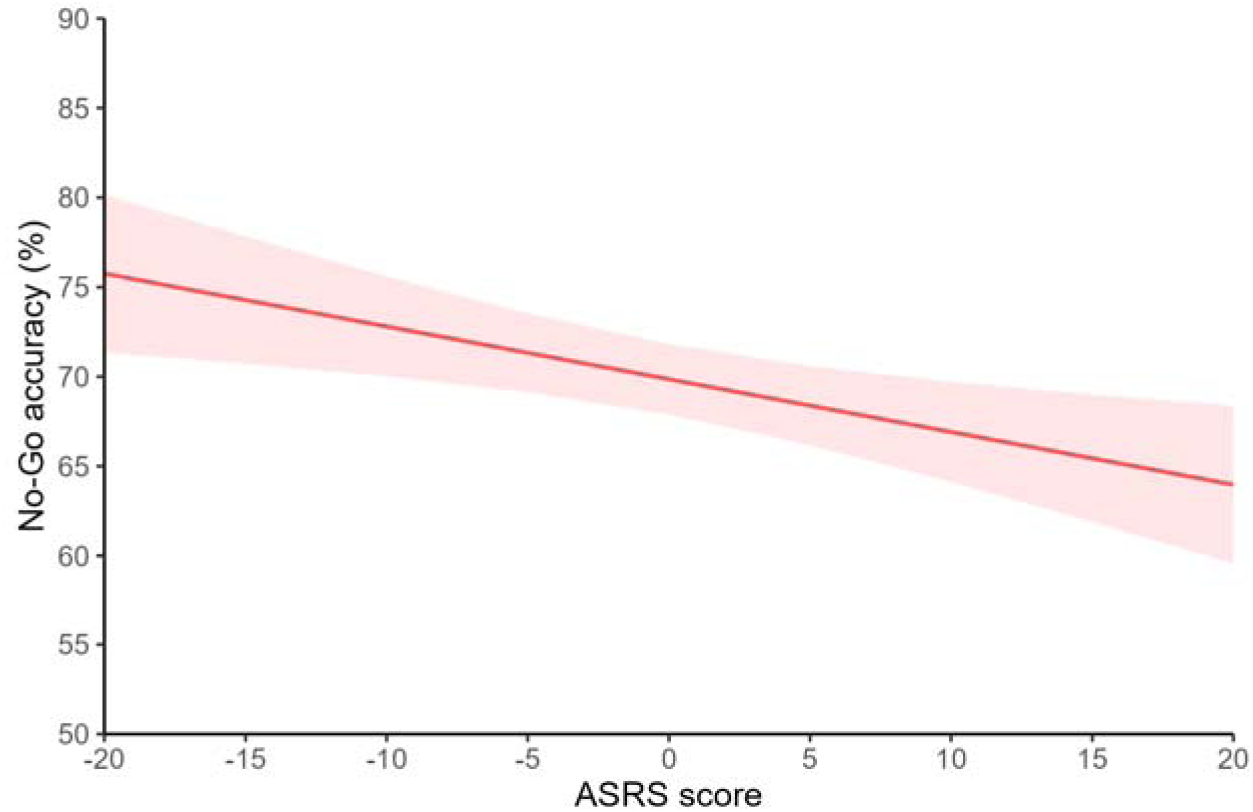
Relationship between ADHD-like symptoms and response inhibition. This figure illustrates the relationship between participants’ self-reported ADHD-like symptoms and their response inhibition. The x-axis represents the ASRS scores, centered around the sample mean, while the y-axis shows No-Go accuracy in percentage. The graph shows a negative relationship, where higher ASRS scores were associated with a decrease in No-Go accuracy. This finding indicates that individuals with more self-reported ADHD-like traits tended to have poorer inhibitory control.

### ADHD-like traits reshape the balance between inhibitory control and statistical learning

We fitted a linear mixed-effects model with the block wise mean reaction time to’go’ trials as the outcome variable. Fixed effects included a four-way interaction between Block (centered), Triplet Type, ASRS scores (centered), and No-Go accuracy (centered). All continuous predictors (Block, ASRS scores, and No-Go accuracy) were mean-centered and treated as scale variables in the model. Random effects included participant ID and random slopes for block within participants, allowing individual variation in learning trajectories across time. The model revealed a significant main effect of Block (*F*_(1, 228.50)_ = 220.16, *p* <.001), indicating that reaction time to’go’ trials significantly decreases across levels of Block (*b* =-1.806, SE = 0.122). The model revealed a significant main effect of Triplet Type (*F*_(1, 13044.25)_ = 83.34, *p* <.001), indicating that reaction time to’go’ trials significantly differed between high and low probability triplets. Participants were faster to high-probability triplets (*b* =-3.067, SE = 0.336), indicating statistical learning. The interaction between Block and Triplet Type was not significant (*F*_(1, 13044.64)_ = 2.57, *p* =.109). The main effect of ASRS was also not significant (*F*_(1, 1224.74)_ = 0.29, *p* =.594), indicating that ADHD-like traits do not influence baseline RT. The main effect of No-Go performance was significant (*F*_(1, 13454.02)_ = 801.40, *p* <.001), indicating faster RT in blocks of weaker No-Go performance (*b* = 55.63, SE = 1.965). This suggests that participants with weaker response inhibition exhibited faster RT. The interaction between No-Go Performance and Block was also significant (*F*_(1, 11556.93)_ = 41.20, *p* <.001). Simple slopes analyses were performed at the beginning, middle, and end points of the Block factor, revealing significant differences between time points in the slopes of No-Go performance (all *p* <.001). The largest difference in RTs based on No-Go accuracy performance was found at the beginning of the task. Participants with better response inhibition (higher No-Go accuracy) showed slower RT, particularly early in the task. This relationship diminished as the task progressed. Post-hoc comparison of estimated marginal trends indicated that the effect of No-Go Performance on’go’ reaction times weakened over time (early blocks: *b* = 67.3, *SE* = 2.82, *p* <.001; middle blocks: *b* = 55.6, *SE* = 1.96, *p* <.001; late blocks: *b* = 43.8, *SE* = 2.54, *p* <.001).

The interaction between No-Go Performance and Triplet Type was also significant (*F*_(1, 13049.04_) = 22.36, *p* <.001). Simple effects analysis confirmed that the difference between high and low probability triplets varied significantly based on No-Go accuracy levels. At low No-Go accuracy, participants demonstrated the largest statistical learning effect (*b* =-9.40, *SE* = 0.979, *p* <.001). This effect remained significant but was diminished at moderate No-Go accuracy (*b* =-6.16, *SE* = 0.672, *p* <.001) and was smallest at high No-Go accuracy (*b* =-2.92, *SE* = 0.940, *p* = 0.002). Pairwise contrasts were conducted to investigate the differences in the statistical learning effect between No-Go accuracy levels. The analysis revealed significant differences across all comparisons. A significant decrease in the learning effect was observed when comparing the low and moderate No-Go accuracy levels (*b* =-3.24, *SE* = 0.685, *p* <.0001). The largest difference was found between the low and high No-Go accuracy levels, where the learning effect was significantly reduced (*b* =-6.48, *SE* = 1.370, *p* <.001). A significant difference was also present between the moderate and high No-Go accuracy levels, with the learning effect being lower in the high accuracy level (*b* =-3.24, *SE* = 0.685, *p* <.001). These results indicate that the magnitude of the statistical learning effect diminishes as No-Go accuracy increases. The three-way interaction between No-Go performance, ASRS score, and Block was also significant (*F*_(1, 12400.85)_ = 8.10, *p* =.004). However, the post hoc test did not reach significance (*p* >.476). The three-way interaction between No-Go performance, ASRS score, and Triplet Type was also significant (*F*_(1, 13046.16)_ = 4.36, *p* =.037). The post hoc test revealed that for participants with low ASRS scores, statistical learning (i.e., the difference between high-and low-probability trials) was gradually worse at higher levels of No-Go performance [low vs. mean: *b* = –4.73, *SE* = 1.03, *p* <.001; low vs. high: *b* = –9.46, *SE* = 2.05, p <.001; mean vs. high: *b* = –4.73, *SE* = 1.03, *p* <.001]. For participants who had mean ASRS scores, statistical learning also gradually declined at higher levels of No-Go performance [low vs. mean: *b* = –3.24, *SE* = 0.69, *p* <.001; low vs. high: *b* = –6.47, *SE* = 1.37, p <.001; mean vs. high: *b* = –3.24, *SE* = 0.69, *p* <.001]. However, at the higher end of the ASRS continuum, this relationship was attenuated and did not reach statistical significance: [low vs. mean: *b* = –1.74, *SE* = 0.95, p = 0.203; low vs. high: *b* = –3.48, *SE* = 1.91, *p* = 0.203; mean vs. high: *b* = –1.74, *SE* = 0.95, *p* = 0.203] (Figure 4). Full results are presented in Supplementary Table 2, including b, SE b, 95% CI, t, df, and p values.

**Figure 4.**
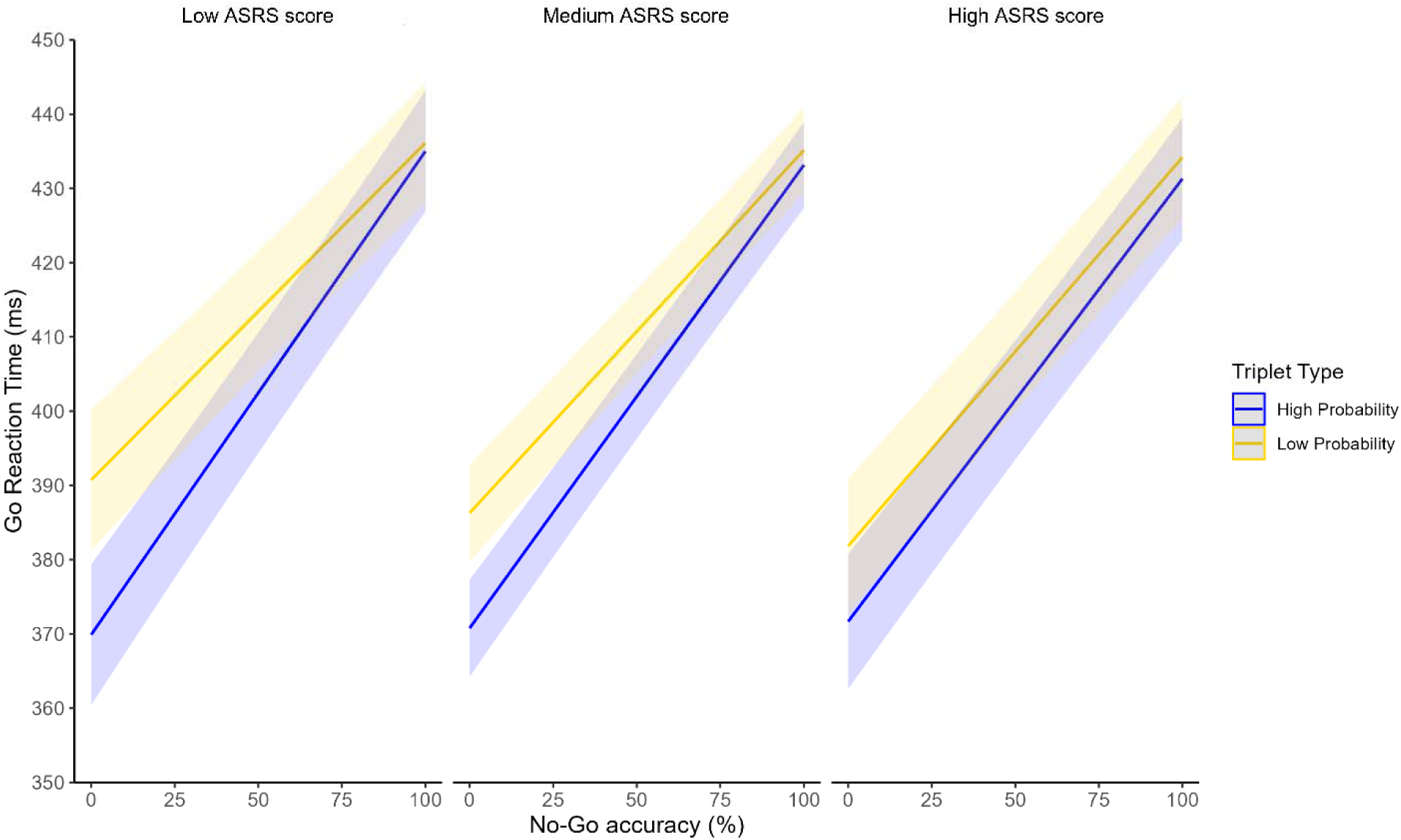
The interaction between response inhibition, ADHD-like symptoms, and statistical learning. This figure illustrates the three-way interaction among No-Go accuracy (as a measure of response inhibition), statistical learning, and ASRS scores (as a measure of ADHD-like symptoms). The x-axis represents No-Go accuracy, while the y-axis shows Go reaction time in ms. The two lines represent reaction times for high-probability triplets (blue line) and low-probability triplets (yellow line). The graph shows how the relationship between response inhibition and statistical learning is moderated by ADHD-like symptom severity. For participants with low or mean ASRS scores, statistical learning declined as No-Go accuracy increased. In contrast, for participants with high ASRS scores, statistical learning remained relatively stable across levels of No-Go accuracy. Please note: For visualization purposes, ASRS scores and No-Go accuracy are plotted at representative values to illustrate the interaction, though both were treated as continuous variables in the statistical models.

In summary, our results revealed that as the task progressed, response inhibition decreased, with participants possessing higher ADHD-like traits exhibiting weaker response inhibition overall. Participants became faster as the task progressed and responded more quickly to high-probability triplets than to low-probability triplets, indicating significant statistical learning. Furthermore, participants with weaker response inhibition exhibited faster RT. Conversely, those with stronger response inhibition showed slower RT, particularly at the beginning of the task. We also observed that the magnitude of the statistical learning effect diminishes as response inhibition increases. Most importantly, unlike individuals with fewer ADHD-related traits, those with a higher symptom load do not show a significant statistical learning advantage even when their response inhibition is poor. Therefore, we conclude that ADHD-related traits alter the dynamic interplay between response inhibition and statistical learning.

## Discussion

Our study reveals how ADHD-like traits shape the dynamic balance between statistical learning and inhibitory control in a large non-clinical sample of young adults. Whereas prior research has typically examined these functions in isolation or in clinical groups, we employed a paradigm that concurrently probed both processes, enabling us to capture their interaction across the full spectrum of ADHD-like traits. We found that higher ADHD-like traits were reliably associated with reduced inhibitory control, confirming a hallmark deficit of ADHD even in individuals below diagnostic thresholds. Critically, we uncovered an antagonistic relationship between inhibition and statistical learning: weaker inhibitory control was linked to enhanced learning of environmental regularities. However, this compensatory advantage was not uniform. Instead, it was progressively attenuated across the ADHD trait continuum and weaker among individuals with high symptom prevalence. These findings provide evidence that ADHD-like traits modulate not only isolated neurocognitive processes but also their interplay, highlighting a spectrum-based mechanism that may underlie the transition from adaptive variability to maladaptive behavior.

Our initial findings, showing that higher ADHD-like symptoms were associated with poorer response inhibition, align with a substantial body of research on the neural underpinnings of ADHD (Cortese, 2012; Cortese et al., 2012; Guo et al., 2020). Deficits in inhibitory control are consistently found in individuals with ADHD (Nigg et al., 2002; Senkowski et al., 2024), while other research has shown that ADHD-like traits can also impair inhibition (Polner et al., 2015). Importantly, we demonstrate that this deficit persists even in a demanding paradigm that simultaneously assesses inhibition and SL, in contrast to previous studies that typically assessed inhibition in isolation (Nigg et al., 2002; Silverstein et al., 2020). This indicates that the inhibitory impairment in ADHD-like traits is robust enough to manifest more real-world-relevant conditions where multiple cognitive systems are engaged in parallel.

While extensive research has examined the relationship between ADHD and various cognitive processes in isolation (Crosbie et al., 2013; Hart et al., 2014; Polner et al., 2015; Wodka et al., 2007), this study addressed a critical gap by investigating their dynamic interaction and how non-clinical ADHD-like traits modulate this. Our findings reveal a strong association between response inhibition and statistical learning, with poorer inhibition corresponding to enhanced statistical learning. Crucially, we revealed that ADHD-like traits significantly influence this relationship, as the learning advantage gradually diminishes across the symptom spectrum and becomes non-significant at the higher end of the ADHD-like trait continuum. These results reinforce the competition hypothesis (Ambrus et al., 2020; Horváth et al., 2022; Janacsek et al., 2012; Nemeth et al., 2013; Pedraza, Farkas, et al., 2024; Pedraza, Vékony, et al., 2024), which posits an antagonistic relationship between executive functions and statistical learning. While individuals with low ADHD-like traits align with this common pattern, the antagonism weakens as symptoms escalate, evolving into an altered form of interaction. This suggests that non-clinical ADHD symptoms play a significant role in shaping the interplay between neurocognitive processes.

This study also highlights the advantages of a spectrum-based methodological approach. Such an approach has already revealed associations with multiple neurocognitive processes in ADHD (Crosbie et al., 2013; Moses et al., 2022; Panagiotidi et al., 2018) and allows for the detection of psychiatric traits before they reach clinical thresholds. This is particularly relevant for ADHD, where diagnostic criteria require that symptoms appear before the age of 12 (American Psychiatric Association, 2022). Adopting a spectrum approach makes it possible not only to identify symptoms that persist from childhood without further causing functional impairment but also to uncover undiagnosed difficulties originating in childhood and to detect adult-onset ADHD (Moffitt et al., 2015).Furthermore, the spectrum approach has proven similarly useful in capturing alterations in the interplay between neurocognitive functions associated with other non-clinical traits (Brezóczki et al., 2025; Colzato et al., 2025). A key question for future research is whether these neurocognitive alterations are trait-or disorder-specific or instead reflect a transdiagnostic general psychopathology factor (Caspi et al., 2014; Gluschkoff et al., 2019)and whether they represent qualitative differences or merely quantitative shifts along the continuum from subclinical to clinical expression.

Our results may also be interpreted through a compelling framework from animal research (Flagel et al., 2007). Researchers examining animal models have found that two distinct phenotypes exist in reward-learning scenarios: sign-trackers and goal-trackers (Felix & Flagel, 2024; Flagel et al., 2007). For sign-trackers, the cue itself becomes so highly valued that it motivates behavior independently of the reward, which can become maladaptive (Flagel et al., 2007). In a classic experimental setup, animals learn that the appearance of a lever predicts the delivery of food. While goal-trackers respond to the lever by moving toward the location where the food will appear, sign-trackers treat the lever itself as the reward, approaching it and even licking or biting it (Felix & Flagel, 2024; Flagel et al., 2007, 2009; Flagel & Robinson, 2017). In these cases, the’logical’ top-down processes that support goal-tracking behavior are overridden by bottom-up urges. This framework provides a plausible explanation for our findings regarding the interplay between inhibition and learning (Felix & Flagel, 2024). In our task, the No-Go stimulus acts as a potent environmental cue. Individuals with lower ADHD-like traits may better manage the’call’ of this stimulus. Even when their inhibition is low, we hypothesize that their other top-down executive functions remain sufficiently intact to regulate behavior. In contrast, participants with high ADHD-like traits may be unable to resist the stimulus itself. For these individuals, the No-Go signal may function as a motivational magnet that triggers an automatic response. Their top-down systems are not robust enough to inhibit the urge to react to the cue, regardless of the task requirements. This explains why the statistical learning’advantage’ seen in other participants breaks down at the higher end of the ADHD symptom continuum.

Another angle of interpretation emerges from research on mind wandering, the spontaneous shift of attention from external demands to internally generated thought (Killingsworth & Gilbert, 2010; Mooneyham & Schooler, 2013; Smallwood & Andrews-Hanna, 2013). Mind wandering, defined as the unintentional shift of attention, has recently been proposed as a potential mechanism underlying ADHD symptoms and associated functional impairments, given its strong correlations with core symptom domains such as inattention (Allan Cheyne et al., 2009; Seli et al., 2015), impulsivity (Allan Cheyne et al., 2009; Arabacı & Parris, 2018), and hyperactivity (Shaw & Giambra, 1993). Research on mind wandering in ADHD has examined both clinical populations and subclinical traits in non-clinical samples, including college students, demonstrating a positive association between mind wandering tendencies and symptom prevalence (Arabacı & Parris, 2018; Biederman et al., 2019; Helfer et al., 2021; Jonkman et al., 2017; Seli et al., 2015). Since attention deficits in ADHD are thought to result from dysregulation in brain networks, with spontaneous streams of thought (mind wandering) emerging as a consequence (Bozhilova et al., 2018; Helfer et al., 2021). This relationship is particularly noteworthy, as mind wandering can influence not only statistical learning (Simor et al., 2025; Vékony, Farkas, et al., 2025), but also the interaction between executive control processes, such as inhibition, and statistical learning (Vékony, Brezóczki, et al., 2025). We can speculate that mind wandering may act as a transdiagnostic factor in ADHD, carrying important implications for both prevention and intervention strategies (Bachmann et.al). On the one hand, it can disrupt executive functioning (Jonkman et al., 2017; Kane et al., 2007); yet in return, it may foster creativity and constructive thought processes (Baird et al., 2012; Mooneyham & Schooler, 2013), and it may also facilitate SL that supports the development of skills and habits (Simor et al., 2025; Vékony, Farkas, et al., 2025). This dual perspective indicates that, although mind wandering entails maladaptive aspects, its adaptive potential could be strategically leveraged in therapeutic approaches to ADHD (Biederman et al., 2017, 2019; Seli et al., 2015), highlighting the need for future research to elucidate how ADHD and mind wandering jointly shape interactions among neurocognitive processes.

## Limitations

Several limitations of the present study should be noted. First, our sample consisted primarily of university students, which resulted in a narrow age range and high educational attainment. While this population is common in cognitive research, it may not be fully representative of the general population. Future studies should include a more diverse age range to determine if the observed interplay between ADHD-like traits and cognitive mechanisms remains consistent across the lifespan. Second, there was a significant gender imbalance in our sample, with 81% identifying as women. Although some research suggests that statistical learning is a robust mechanism that does not differ significantly between sexes (Botia et al. 2025), findings in the field of response inhibition are less conclusive (Bjorklund et al., 1996; Erickson et al., 2005; MacApagal et al., 2011; Sjoberg & Cole, 2018; Thakkar et al., 2014; Wright et al., 2014). Therefore, our results should be interpreted with caution regarding their generalizability to men, and future research should aim for a more balanced gender distribution to investigate potential sex-specific cognitive strategies.

Furthermore, the entire experimental protocol was conducted online without direct supervision. While we applied rigorous exclusion criteria to minimize the impact of uncontrolled variance, these factors cannot be entirely eliminated (Rodd, 2024). Consequently, these unsupervised conditions may have influenced the strength and stability of the observed relationship between inhibitory control and statistical learning across the ADHD spectrum. While online testing allowed for a broader reach, future research conducted in controlled laboratory settings would be beneficial to replicate these findings and further validate the observed antagonistic effects.

Another limitation concerns the exclusive reliance on the ASRS (Kessler et al., 2005; v1.1) for estimating ADHD-like traits. While self-report questionnaires are efficient and widely validated for dimensional studies in the general population, they may lack the precision of multi-instrument or multi-informant approaches. Factors such as social desirability or limited self-awareness could influence the estimation of trait-related gradients along the ADHD continuum. Future studies would benefit from incorporating collateral reports from family members or additional validated scales such as the Conners’ Adult ADHD Rating Scales (CAARS; Conners, Erhardt, & Sparrow, 1999) or the Wender Utah Rating Scale (WURS; Ward et al., 1993). Combining these measures would provide a more robust characterization of ADHD-like symptoms and their relationship with cognitive performance.

### Future directions

To build on the current findings, future research should use a wider variety of tests to better understand the relationship between ADHD-like traits and cognitive processes. A logical next step involves incorporating additional diagnostic and symptom-based measures, such as the Conners’ Adult ADHD Rating Scales (CAARS) or the Wender Utah Rating Scale (WURS) (Macey, 2003; Ward et al., 1993). These tools would allow researchers to more precisely evaluate ADHD symptoms, including those from childhood, and provide a richer context for the current findings. To develop a more comprehensive cognitive profile, future studies should also measure other core executive functions commonly affected by ADHD, including working memory and cognitive flexibility. Investigating the interplay among these functions and their relationship to statistical learning and inhibitory control is a crucial next step. Indeed, we propose that the behavioral manifestations of ADHD may not stem from isolated impairments but rather from a disruption in the dynamic interaction among different cognitive systems. Future research should systematically examine this interplay to fully elucidate the underlying mechanisms of ADHD symptoms and how they contribute to maladaptive behavioral patterns. Additionally, by comparing individuals with subclinical traits to those with formal clinical diagnosis, we can gain a more comprehensive understanding of how these cognitive functions differ across the entire spectrum of symptomatic expression. Our study highlights the need for this approach by confirming that inhibitory deficits, as well as shifts in the dynamic interaction between inhibition and learning, are present even in a non-clinical sample. Because these neurocognitive changes emerge before symptoms reach a clinically significant threshold, assessing both individual functions and their interplay across the full ADHD spectrum is crucial for early detection. Finally, the theoretical framework proposed by our research has broader implications beyond ADHD. Future research could explore whether this trait-based approach to cognitive function applies to other disorders that show high comorbidity with ADHD. This would be a crucial step toward developing a new theoretical framework for diagnosis and intervention that moves beyond traditional diagnostic boundaries and is better equipped to support individuals with varying levels of symptomatic severity.

## Conclusion

Our study demonstrates that ADHD-like traits influence not only inhibitory control and learning in isolation but also their dynamic interaction, revealing a cognitive trade-off that is systematically altered across the symptom spectrum. This finding underscores the importance of moving beyond binary case–control frameworks toward dimensional models that capture variability in the general population. Cognitive alterations can emerge even in individuals below diagnostic thresholds, with implications for everyday functioning and vulnerability to maladaptive outcomes. Recognizing ADHD as a spectrum condition opens new opportunities for trait-based approaches that emphasize early detection and proactive intervention. Rather than waiting for symptoms to escalate into a formal diagnosis, clinicians could leverage these insights to design targeted strategies that strengthen both executive control and learning processes, as well as their interaction. Such an approach would broaden access to support, reduce stigma, and facilitate personalized psychoeducation. More broadly, our findings highlight the need for a new theoretical framework that situates ADHD within a continuum of cognitive functioning, with direct relevance for transdiagnostic psychiatry and precision mental health.

## Acknowledgement

This work was supported by the French National Research Agency (ANR-24-CE37-5807), the National Brain Research Program (NAP2022-I-2/2022), and the Hungarian Scientific Research Fund (NKFIH 153150) all awarded to D.N.; and the EKÖP-24 University Excellence Scholarship Program of the Ministry for Culture and Innovation from the Source of the National Research, Development and Innovation Fund (EKÖP-24-3-II-ELTE-1159 (to B.B); the Spanish Ministry of Science, Innovation and Universities (MICIU), the State Research Agency (AEI), and the European Regional Development Fund (FEDER, UE) through the grant PID2024-160183NA-I00 (MICIU/AEI/10.13039/501100011033/FEDER, UE) (to T.V.).

## Contributions

K.H.: Data Curation, Formal analysis, Writing – Original Draft, Writing – Review & Editing, Visualization; B.B.: Conceptualization, Software, Validation, Investigation, Data Curation, Formal analysis, Writing – Original Draft, Writing – Review & Editing, Funding; A.H.: Formal analysis, Writing – Review & Editing, T.V.: Conceptualization, Software, Validation, Investigation, Data Curation, Formal analysis, Visualization, Resources, Writing – Review & Editing, Supervision; D.N.: Conceptualization, Software, Validation, Resources, Writing – Review & Editing, Supervision, Funding.

## Data availability statement

Data and codes for analysis are available on the following link: https://osf.io/evm2k/

