## Supplementary information for "ADHD-like traits reshape the balance between inhibitory control and predictive processes" (from another version of this preprint: version 1, 2025-10-29)

**Supplementary Material**

**Details of the models presented in the main text**

**Supplementary Table 1. The relationship between ADHD-like traits and response inhibition.**

|  | **No-Go Accuracy** | | | | | |
| --- | --- | --- | --- | --- | --- | --- |
| *Terms* | *b* | *SE b* | *95% CI* | *t* | *df* | *p* |
| (Intercept) | 0.6985 | 0.0101 | 0.6787 – 0.7184 | 69.3458 | 223.9624 | **<0.001** |
| Block | -0.0064 | 0.0005 | -0.0075 – -0.0054 | -12.3736 | 224.6086 | **<0.001** |
| ASRS score | -0.0029 | 0.0010 | -0.0049 – -0.0010 | -2.9303 | 223.9363 | **0.004** |
| Block × ASRS score | -0.0000 | 0.0001 | -0.0001 – 0.0001 | -0.6609 | 224.0943 | 0.509 |
| **Random Effects** | | | | | | |
| σ^2^ | 0.08 | | | | | |
| τ_00_ _ID_ | 0.02 | | | | | |
| τ_11_ _ID.Block_ | 0.00 | | | | | |
| ρ_01_ _ID_ | 0.21 | | | | | |
| ICC | 0.24 | | | | | |
| N _ID_ | 226 | | | | | |
| Observations | 11965 | | | | | |
| Marginal R^2^ / Conditional R^2^ | 0.038 / 0.272 | | | | | |

*Note.* Response inhibition was weaker among ADHD-like traits (main effect of ASRS score), and response inhibition became weaker as the task progressed (main effect of Block).

**Supplementary Table 2. The interplay between statistical learning and response inhibition among ADHD-like traits**

|  | **Reaction Time** | | | | | |
| --- | --- | --- | --- | --- | --- | --- |
| *Terms* | *b* | *SE b* | *95% CI* | *t* | *df* | *p* |
| (Intercept) | 417.1313 | 2.8443 | 411.5263 – 422.7363 | 146.6535 | 224.5327 | **<0.001** |
| Block | -1.8056 | 0.1217 | -2.0453 – -1.5658 | -14.8378 | 228.5000 | **<0.001** |
| Triplet Type [High] | -3.0675 | 0.3360 | -3.7261 – -2.4089 | -9.1289 | 13044.2540 | **<0.001** |
| ASRS score | -0.1518 | 0.2840 | -0.7115 – 0.4079 | -0.5345 | 224.7405 | 0.594 |
| No-Go Accuracy | 55.6279 | 1.9650 | 51.7762 – 59.4797 | 28.3091 | 13454.0158 | **<0.001** |
| Block ×Triplet Type [High] | -0.0626 | 0.0390 | -0.1390 – 0.0139 | -1.6046 | 13044.6411 | 0.109 |
| Block ×ASRS score | 0.0088 | 0.0122 | -0.0152 – 0.0328 | 0.7213 | 230.1420 | 0.471 |
| Triplet Type [High] ×ASRS score | 0.0508 | 0.0340 | -0.0158 – 0.1174 | 1.4963 | 13044.2629 | 0.135 |
| Block × No-Go Accuracy | -1.3342 | 0.2078 | -1.7416 – -0.9267 | -6.4190 | 11556.9297 | **<0.001** |
| Triplet Type [High] ×No-Go Accuracy | 6.7464 | 1.4268 | 3.9497 – 9.5431 | 4.7284 | 13049.0377 | **<0.001** |
| ASRS score × No-Go Accuracy | 0.0365 | 0.1932 | -0.3422 – 0.4153 | 0.1891 | 13409.0745 | 0.850 |
| (Block × Triplet Type [High] × ASRS score | -0.0040 | 0.0040 | -0.0117 – 0.0038 | -1.0105 | 13044.4802 | 0.312 |
| (Block × Triplet Type [High] × No-Go Accuracy | 0.2929 | 0.1604 | -0.0216 – 0.6074 | 1.8256 | 13052.3196 | 0.068 |
| (Block × ASRS score × No-Go Accuracy | 0.0595 | 0.0209 | 0.0185 – 0.1005 | 2.8464 | 12400.8549 | **0.004** |
| (Triplet Type [High] ×ASRS score × No-Go Accuracy | -0.3103 | 0.1486 | -0.6017 – -0.0190 | -2.0880 | 13046.1600 | **0.037** |
| (Block × Triplet Type [High] × ASRS score × No-Go Accuracy | -0.0039 | 0.0169 | -0.0370 – 0.0291 | -0.2341 | 13047.7679 | 0.815 |
| **Random Effects** | | | | | | |
| σ^2^ | 1411.73 | | | | | |
| τ_00_ _ID_ | 1801.51 | | | | | |
| τ_11_ _ID.Block_ | 2.98 | | | | | |
| ρ_01_ _ID_ | -0.30 | | | | | |
| ICC | 0.59 | | | | | |
| N _ID_ | 226 | | | | | |
| Observations | 13514 | | | | | |
| Marginal R^2^ / Conditional R^2^ | 0.140 / 0.647 | | | | | |

*Note.* Worse No-Go performance was associated with faster RTs (main effect of No-Go). Statistical learning was better during periods of weak No-Go performance (interaction between No-Go and Triplet Type). When No-Go performance was weak, participants showed greater statistical learning (interaction between No-Go and Triplet Type). The significant Triplet Type × No-Go Accuracy × ASRS score interaction revealed that ADHD-like traits modulate this antagonistic relationship.
